# Neural correlates of online action preparation

**DOI:** 10.1101/2023.06.19.545487

**Authors:** Mahdiyar Shahbazi, Giacomo Ariani, Mehrdad Kashefi, J. Andrew Pruszynski, Jörn Diedrichsen

## Abstract

When performing movements in rapid succession, the brain needs to coordinate preparation of an upcoming action with ongoing execution. Here we identify the processes and brain areas involved in this ability. Human participants performed pairs of single-finger presses or three-finger chords in settings where they had to prepare the second movement before or after the first response. Despite matched perceptual and movement requirements, 7T functional MRI revealed increased brain activity in regions along the intra-parietal sulcus and ventral visual stream when actions overlapped. Multivariate analyses suggested that these areas were involved in stimulus identification and action selection. In contrast, the dorsal premotor cortex, known to be involved in planning upcoming movements, showed no discernible signs of heightened activity. This observation suggests that the bottleneck during simultaneous action control and preparation arises at the level of stimulus identification and action selection, whereas movement planning in the premotor cortex can unfold concurrently with execution without interference between the two processes.

**Summary:** The brain’s ability to prepare for upcoming actions while controlling ongoing movements is a crucial evolutionary adaptation of the action system. However, the neural basis of online action preparation remains largely unknown. We found that superior-parietal and occipito-temporal areas exhibited heightened activation during online preparation. Surprisingly, the dorsal premotor cortex, known to be a crucial structure in motor planning, did not display additional activation for overlapping actions. These findings imply that while motor planning within the premotor cortex can occur in parallel with the execution of ongoing movement, the parallel stimulus identification and action selection in the posterior parietal cortex requires additional neural processes.

**Highlights:**

- Individuals prepare upcoming actions while simultaneously controlling ongoing movement
- When these two processes overlap, superior-parietal and occipito-temporal areas show increased activation
- Multivariate analysis suggests that increased activation arises to resolve simultaneous stimulus identification and action selection
- The premotor cortex, known to be involved in motor planning, coordinates simultaneous planning and execution without extra neural engagement

## Introduction

Most laboratory studies of perception, action, or decision-making consist of distinct trials: a sequence of stimulus, response, and feedback is followed by a short pause before the next trial begins. In contrast, most tasks in real life involve a fluid and continuous stream of actions. For example, imagine you are trying to play a piece of music on a piano, aiming to press the right keys with the correct timing. One intuitive strategy is to read only the upcoming chord from the sheet of music, prepare the appropriate hand configuration, and then play the chord. However, this strategy fails if there is not enough time to prepare the next chord after playing the current one. In such situations, one needs to expand their window of preparation by starting to prepare the next movements before the current movement is completed. A series of recent studies show that participants indeed engage in this process of *online preparation* (Ariani & Diedrichsen, 2019; Ariani et al., 2021, 2020; Kashefi et al., 2023), preparing 2-3 movements ahead of the current action. However, which processes and brain areas are involved in coordinating the preparation for future movements with the simultaneous execution of ongoing movement remains unknown.

Electrophysiological recordings in primary motor and premotor regions have shown that the same areas and often even the same neurons are engaged in the preparation and subsequent execution of a single movement (Ames et al., 2014; Ariani et al., 2018, 2022; Crammond & Kalaska, 1994, 2000; Tanji & Evarts, 1976). Given this overlap, how does the brain avoid interference when the execution of one movement occurs simultaneously with the preparation of another? One possibility is that coordinating simultaneous preparation and execution may require increased brain resources, either by recruiting more neurons in the same brain regions or by engaging new areas that were previously not involved in the task. Alternatively, given the evolutionary importance of acting quickly, the brain may have evolved efficient mechanisms to orchestrate simultaneous preparation and execution within the same areas and without interference. This view is compatible with findings from recordings in the primary motor and premotor cortex of non-human primates, showing that movement preparation and execution take place in distinct neural state spaces (Kaufman et al., 2014; Zimnik & Churchland, 2021).

Here we test these two hypotheses across the entire human visuomotor system, which supports a complex cascade of processes during action preparation: First, an external stimulus must be identified, and the next action goal needs to the selected among a set of alternatives (Rosenbaum & Kornblum, 1982). After goal selection, the motoric details of the upcoming action need to be specified. This process brings the motor system to the optimal initial state for the execution (Shenoy et al., 2013). Interference between the current and a future movement may arise at any of these stages of action preparation.

In the current study, we aimed to pinpoint at which processing stage and in which neural regions additional neural processes are required for online preparation. We designed a high-field (7T) fMRI experiment in which participants performed series of finger presses in response to arbitrary visual cues. In the “overlap” condition, preparation for the next movement overlapped with the execution of the ongoing movement. In the “non-overlap” condition, movement preparation and execution occurred sequentially. Since overlap and non-overlap conditions were matched in basic perceptual processes and execution requirements, their difference highlighted the areas that were more activated during online preparation, which could be related to selection, planning, or both. To distinguish between these different processes, we also varied the motoric complexity of the movements from single finger press to a chord of three finger presses. In Experiment 1, comparing these different levels of action complexity while matching cue identification and action selection informed us whether motor planning takes longer for chords than single finger presses. Based on these results, in Experiment 2, any neural difference between the chord and single finger conditions could be attributed to the increased demands in motor planning or execution. Finally, counterbalancing stimulus-response mapping across participants allowed us to use multivariate analysis techniques to distinguish between areas showing preferential processing of cues or actions.

## Results

### Complex actions require longer preparation time

In Experiment 1, we instructed participants (N=11; 4 female; age=26±4) to use their right hand to press a single key or to simultaneously press three keys (chord) on a piano-like keyboard. In both the single finger and chord conditions, 5 different symbols indicated one of 5 actions (Figure 1A). Participants were presented with a sequence of alternating high and low-pitch tones (black and gray notes, respectively, Figure 1B). They were instructed to produce an action in synchrony with the high-pitch tones, according to the symbol that appeared at a variable interval (240-1750 ms) before that tone. We counterbalanced symbols across participants to control for the impact of specific cues on the preparation time, such that any difference in preparation time between the conditions could be attributed to the increased demands on motor planning for the motorically more difficult chords.

**Figure 1.**
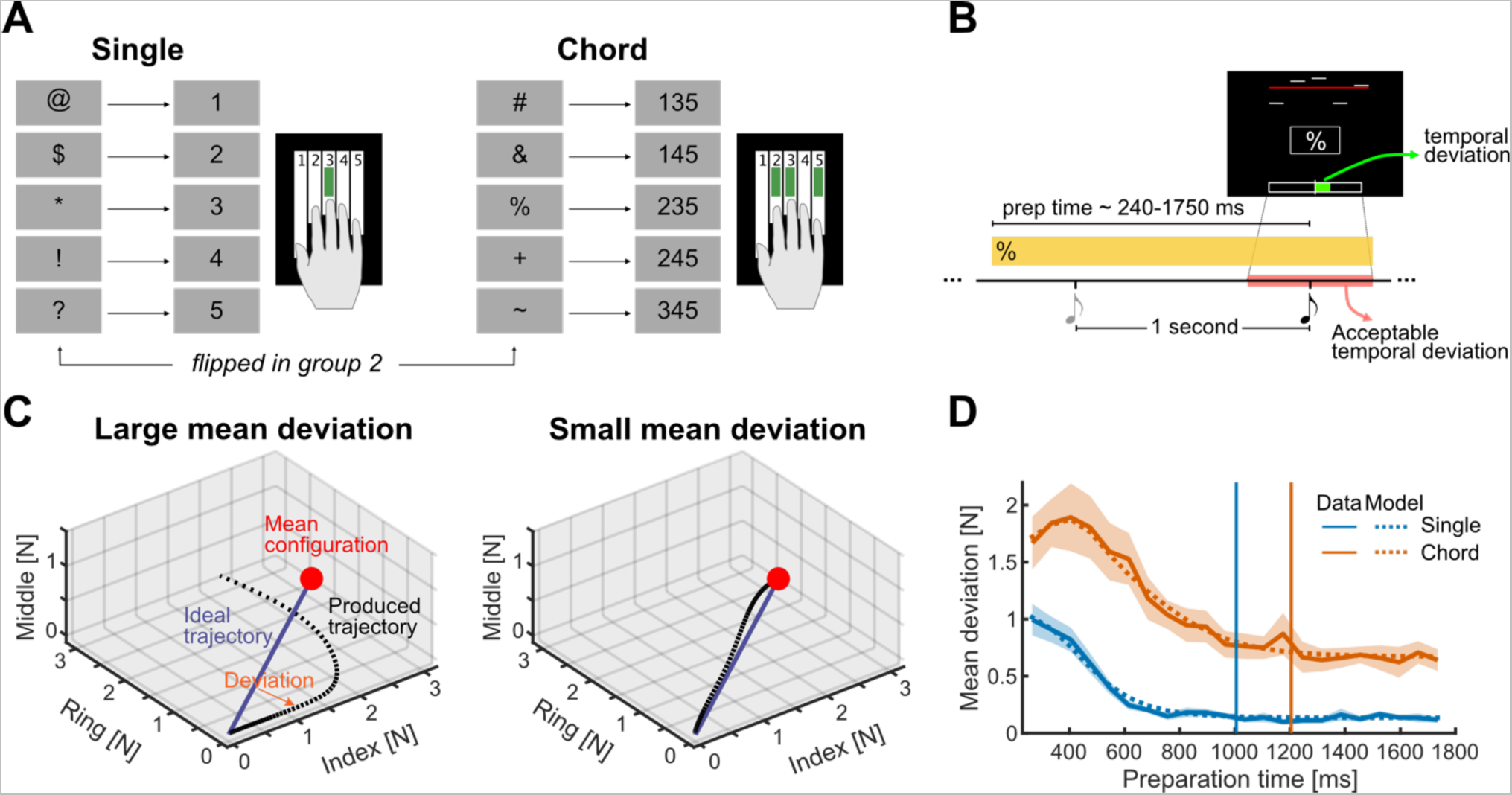
Three-finger chords take longer to prepare. (**A**) Single-finger presses and chords. Participants were divided into two groups, which practiced the same presses, but the symbol-response associations were switched across press types. (**B**) Participants needed to produce presses indicated in the box simultaneously with high-pitch tones (black notes). The presence of the symbol on the screen is represented by the yellow strip. The response window was ±200m (red area), and deviations from the ideal timing were indicated by the bar at the bottom of the screen. For incorrect presses, the bar turned red. Five small lines on the top represented the applied force on each key, with a press threshold of 0.8 N indicated by the red line. (**C**) Example of two trials with produced force trajectories (black dotted lines) with either large (left) or small deviations (right) from the ideal force trajectory (purple line). Trials are illustrated in a three-dimensional finger force space consisting of two active fingers (index and ring) and one passive digit (middle). (**D**) Solid lines indicate the group-averaged mean deviation as a function of available preparation time for single (blue) and chord (orange). The dotted lines are the model fit average across subjects. The vertical lines indicate the average time when the fitted curves for chord (orange) or single finger (blue) reach 105% of the asymptote performance.

When there was not enough time to prepare an action, participants did not manage to press the required fingers accurately and simultaneously. Sequential presses, or initial errors that were later corrected, led to a deviation of their force trajectory from the ideal straight-line trajectory to the desired goal force (Figure 1C). This average deviation (see Methods) decreased with increasing preparation time (Figure 1D). However, even for long preparation times, chords remained more difficult than single-finger presses. To evaluate the required preparation time for both, we fit a Gaussian function with a constant offset (see Methods) to the data of each press type. Based on these fits, we estimated the required preparation time for each participant as the time required for their mean deviation to get within 105% of their performance with full preparation. The required preparation time was 200±47 ms (*t_10_=4.21, p=0.0018*) longer for chord (1204±75ms) than for single finger presses (1005±51ms). These results demonstrate that it takes longer for chords to be fully prepared than single-finger presses even when the requirements of cue identification and action selection are matched.

### Online preparation improves execution quality

In the main experiment (Experiment 2), we aimed to create a condition in which participants (N=22; 12 female; age=24±4) would have to prepare for the next action while still executing the previous one. To do so, we selected a time between two consecutive responses that was not enough to fully prepare a chord and barely enough to prepare a single finger press. Based on the results from Experiment 1, we chose an interval of 750ms. We reasoned that if the stimulus for the second action was presented before the first response was completed, participants would utilize that information to improve their execution quality.

In this experiment, participants were played a series of three sounds consisting of one low-pitch tone followed by two high-pitch tones. They were also shown two boxes on the screen. Symbols appeared first in the upper box and then moved to the lower box 550 ms before each high-pitch tone (Figure 2A-C). Simultaneously with each high-pitch tone, participants had to produce the action indicated by the symbol in the lower box.

**Figure 2.**
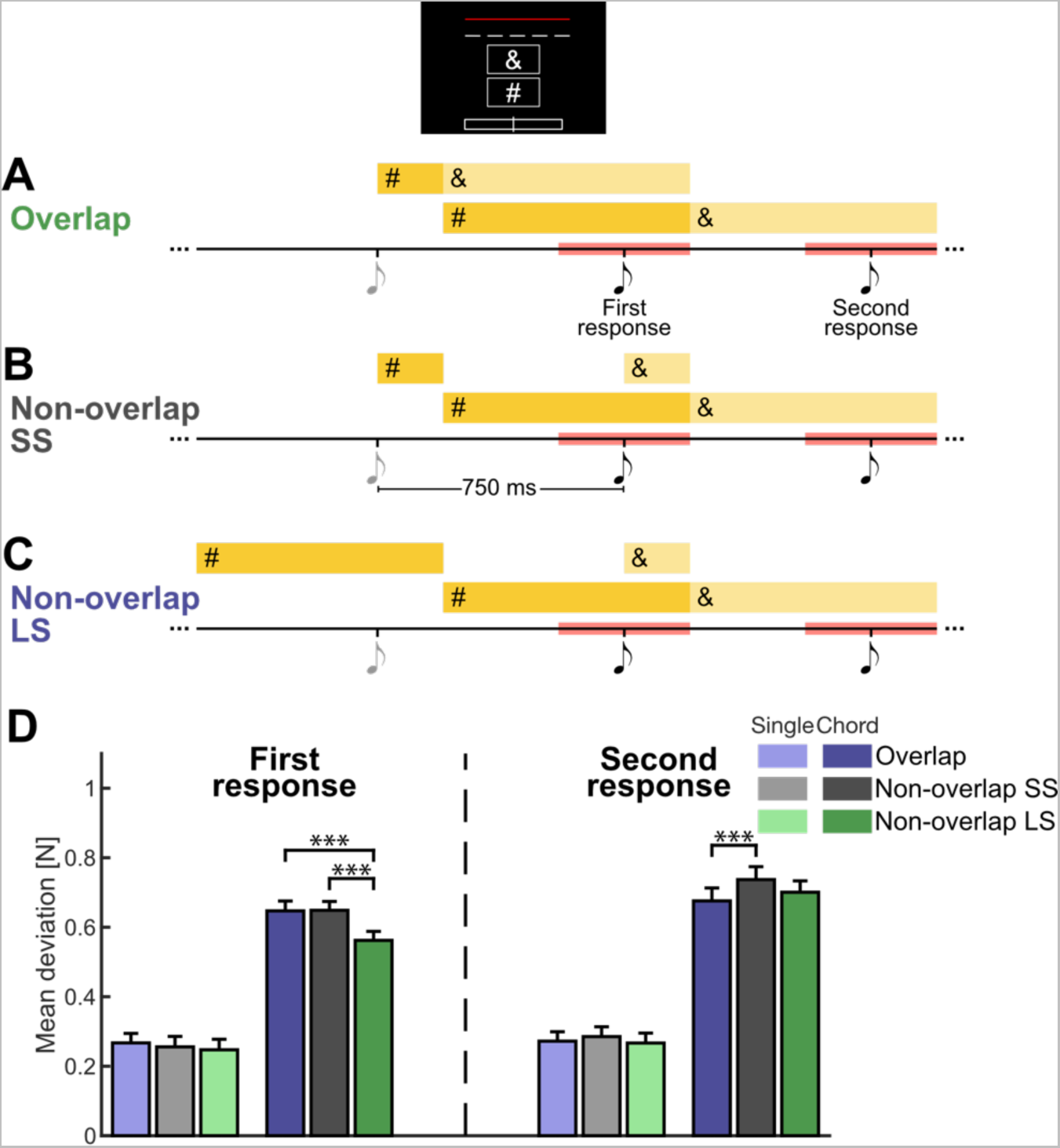
Task design for experiment 2. (**A**) In a continuous paradigm, one low-pitch tone (gray note) alternated with two high-pitch tones (black note). Participants had to produce actions indicated in the lower box synchronously with high-pitch tones with ±200ms (red intervals). The content of the upper box over time is indicated by the upper yellow strips, while the lower yellow strips represent the content of the lower box over time. Distinct shades of yellow were used to differentiate between the time to prepare the first response and the second response. In the overlap condition, the second symbol (&) was visible in the upper box during the preparation phase of the first response. The inset shows the screen during this overlapping preparation period. (**B**) In the non-overlap short-short (SS) condition, the second response cue appeared together with the first tone, leaving participants with no chance to do online preparation. The short preparation time for the first response allowed for a strong test of the effect of online preparation on behavior. (**C**) Same as B, with the difference that the first response in the non-overlap long-short (LS) condition had a long preparation time. This allowed us to compare the fMRI activity averaged across the two conditions to the overlap condition. (**D**) Mean deviation from a straight-line force trajectory (execution quality) for the first response (left) and second response (right) separate for single finger and chords and overlap and non-overlap conditions. Error bars indicate the SEM across participants. ***p<0.001, two-tailed paired-samples t-test.

In two conditions, the response preparation and execution of the two actions occurred sequentially – the symbol indicating the second response (the & in Figure 2) was presented at the exact moment of the tone for the first response. The two non-overlap conditions only differed by the preparation time for the first response. In the short-short condition (SS, Figure 2B), the first stimulus appeared 750 ms before the first high-pitch tone (200 ms in the upper, 550 ms in the lower box). In the long-short condition (LS, Figure 2C), the first stimulus appeared 1300 ms before the first high-pitch tone.

These two conditions were compared to the overlap condition (Figure 2A), where the symbol for the second response was on the screen even 550 ms before the first response execution, resulting in a total preparation time of 1300 ms for the second action. The first response had a short preparation time (750ms). The overlap condition, therefore, only differed from the non-overlap SS condition by the earlier presentation for the second response, allowing for a strong comparison of the response quality for the second response. The non-overlap LS was designed to have an equal average preparation time for response as the overlap condition. This condition was, therefore, ideal for the comparison of activations between conditions, given the limited temporal resolution for fMRI forced us to average activity across each pair of responses.

As intended, the equal preparation time in overlap and non-overlap SS conditions resulted in similar execution quality for the first response (chord: *t_21_=-0.100, p=0.9214*, single: *t_21_=1.481, p=0.1536*). This assured us that the earlier appearance of the second stimulus in the overlap condition did not interfere with the first action. Naturally, the non-overlap LS condition showed a lower mean deviation for the first chord as compared to both non-overlap SS (*t_21_=4.842, p=9e-5*) and overlap conditions (*t_21_=3.677, p=0.0014*), reflecting the longer preparation time. We also observed a similar pattern of results for single-finger conditions. However, the benefit of a longer preparation time was smaller for single finger press compared with chord, as shown by a significant condition x press type interaction (*F_1,21_=8.091, p=0.0011*).

Critically, we found that the participants used the earlier appearance of the first cue to plan the second response during the first response preparation and execution. For chords, the mean deviation for the second response was statistically smaller than in the relevant control condition (non-overlap SS, *t_21_=-5.125, p=4e-5*). A similar pattern also emerged for single finger presses, even though the differences between overlap and non-overlap SS conditions were not significant (*t_21_=-1.505, p=0.1473*). Although we cannot rule out that online preparation provided a small advantage even for single-finger presses, the effect was certainly smaller than for chords, as shown by a significant condition x press type interaction (*F_1,21_=4.387, p=0.0186*). Taken together, our results show that preparation during the ongoing movement can improve the execution quality of the upcoming response, with the benefit becoming more apparent for more complex actions.

### Online preparation activates the superior-parietal lobule and ventral visual stream but not the premotor areas

To examine which brain regions are more activated in online preparation, we compared the fMRI activation during the overlap vs. the non-overlap conditions. Because it is challenging to separate the fMRI activity related to the first and second response, we matched the average preparation time across the two presses between the overlap (Figure 2A) with the non-overlap LS condition (Figure 2C). With average preparation times and execution requirement matched, any difference between the two conditions should be associated with the fact, that in the overlap condition, the second response was prepared in parallel with the first action.

Averaged across single fingers and chords and compared to rest, the task activated the primary and secondary sensorimotor cortices, regions along the intra-parietal sulcus, areas in the occipito-temporal cortex, and the auditory cortex (Figure 3B). Within this task-relevant network, we found that only areas in the superior posterior lobule (SPLa: *t_21_=3.298, p=0.0034*, SPLp: *t_21_=3.269, p=0.0036*) and in the ventral visual stream (MT+: *t_21_=8.044, p=7e-08*, VSVC: *t_21_=6.306, p=3e-06*) were more activated during online preparation (Figure 3C). This was also clear in the surface-based analysis, in which the largest significant clusters (*p=8e-07*, corrected for multiple comparisons, see Methods) extended from the intra-parietal sulcus to visual areas on the boundary between occipital and temporal lobe (dashed back line, Figure 3C).

**Figure 3.**
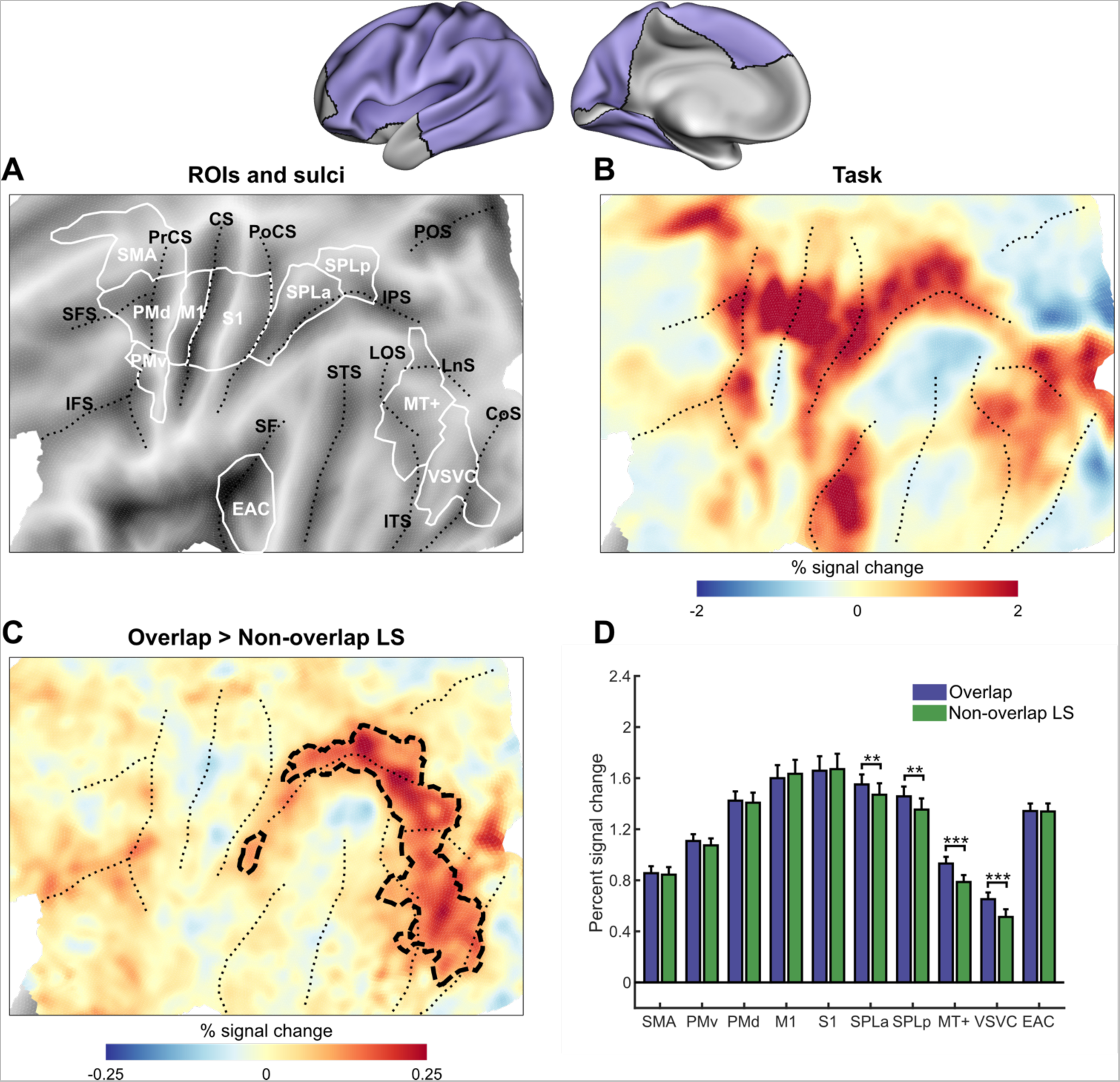
Online preparation activates superior-parietal and occipito-temporal regions. The inset shows the inflated cortical surface of the contralateral (left) hemisphere, highlighting the area of interest (A-C, purple). **(A)** Flat representation of the neocortex with major sulci indicated by black dotted lines and regions of interest by white borders. (**B**) Group-averaged percent signal change during task vs. resting baseline averaged across overlap and non-overlap conditions and press types (single, chord). (**C**) The difference in percent signal change between overlap and non-overlap LS conditions averaged across single finger and chord. Black dashed boundaries represent significant clusters. (**D**) Average percent signal change in the predefined ROIs for overlap (purple) and non-overlap LS (green) conditions. Error bars indicate SEM across participants. ROIs: early auditory cortex (EAC), ventral stream visual cortex (VSVC), MT+ complex and neighboring visual areas (MT+), posterior superior-parietal lobule (SPLp), anterior superior-parietal lobule (SPLa), primary somatosensory cortex (S1), primary motor cortex (M1), dorsal premotor cortex (PMd), ventral premotor cortex (PMv), secondary motor area (SMA). Sulci: superior frontal sulcus (SFS), inferior frontal sulcus (IFS), precentral sulcus (PrCS), central sulcus (CS), post central sulcus (PoCS), intra-parietal sulcus (IPS), parieto-occipital sulcus (POS), lateral occipital sulcus (LOS), lunate sulcus (LnS), superior temporal sulcus (STS), inferior temporal sulcus (ITS), collateral sulcus (CoS), sylvian fissure (SF). Significant pairwise differences are indicated with **p<0.005, ***p<0.001, two-tailed paired-samples t-test.

Interestingly, we did not find any evidence of extra activation during online preparation in premotor areas. Neither the dorsal premotor cortex (PMd, *t_21_=0.898, p=0.3791*) nor the supplementary motor area (SMA, *t_21_=0.560, p=0.5812*) was significantly more active during the overlap condition. This is despite the fact that it is well established that both regions play an important role in motor planning of hand and arm movements (Gallivan et al., 2016, 2011; Henderson et al., 2022; Hoshi & Tanji, 2004; Shenoy et al., 2013; Tanji & Shima, 1994). These results demonstrate an important dissociation, with posterior parietal, but not frontal premotor areas, showing more activation during online preparation.

### Functional role of online preparation area

We then asked which processes (cue identification, action selection, motor planning, and motor execution) occurred in the areas that were more highly activated during the overlap condition. We performed two analyses to obtain insight into this question.

First, we compared the fMRI activation for single finger press vs. chords, two actions that differed in motoric complexity with equivalent visual stimuli and number of alternative options. As expected, we found that M1 and S1 (M1: *t_21_=10.258, p=1e-09*, S1: *t_21_=10.41, p=9e-10*) were more activated during the chord than the single finger conditions. Additionally, we observed that pressing chords elicited more activity in premotor cortical areas (PMd: *t_21_=8.979, p=1e-08*, PMv: *t_21_=6.163, p=4e-06*, SMA: *t_21_=7.290, p=3e-07*) and superior-parietal lobule (SPLa: *t_21_=9.217, p=7e-09*, SPLp: *t_21_=6.776, p=1e-06*). We did not find evidence for higher activation for complex actions in auditory areas (EAC: *t_21_=1.601, p=0.1243*). Surprisingly, given that chord and single-finger conditions were matched in terms of the visual information, we found that activity in the lateral occipito-temporal cortex was modulated by the complexity of the motor response (MT+: *t_21_=5.650, p=1e-05*, VSVC: *t_21_=5.520, p=2e-05*).

Second, we used multi-voxel pattern analysis to characterize the features that are represented in the fine-grained patterns of activity in these regions. For this, we required an estimate of the activity patterns for the 10 possible actions (5 different singles and 5 chords) separately. This was done by modelling the first and second responses separately and averaging across all responses and conditions (see Methods). We then asked to what degree the similarity between activity patterns can be explained by the similarity between visual cues or the similarity between actions. As a model, we used the data from the other half of the participants, for which the assignment between visual cues to actions was switched. For example, to assess the similarity of patterns within SPLa, we used the average similarities among actions within SPLa of the other half of the participants as a model for action-related processes. Additionally, we employed the similarity between the cues measured at the same time as a model for cue-related processes. To be able to model combinations of both cue- and action-related processes, we used a Bayesian model family approach (Yokoi & Diedrichsen, 2019), which evaluates the evidence for each model component (cue or action) in the context of the other component.

In primary motor and visual areas, the resultant subject-averaged log Bayes-factor for action and cue encoding provided the expected results. Motor areas, including M1, S1, and PMd, showed strong action encoding (Figure 4C), whereas high-level visual areas represented the visual cues (Figure 4D). In these areas, we found no evidence for action-related encoding, even though these regions showed higher activity during the chord than single finger conditions. This suggests that even visual processing needs to work harder or longer if motor planning takes longer (see Discussion).

**Figure 4.**
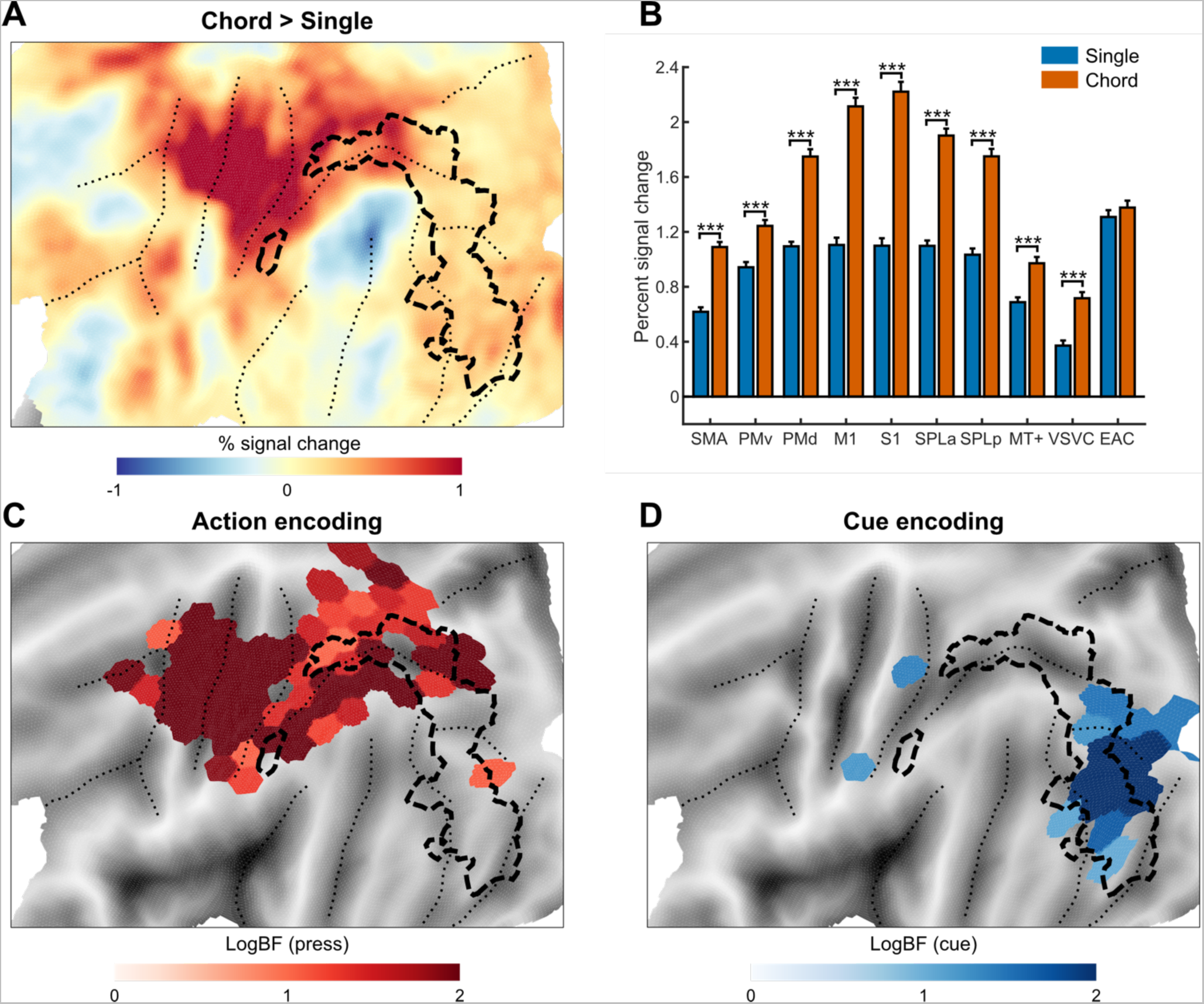
Evidence for involvement in cue-related and action-related processes. (**A**) The difference in evoked activation between chord and single-finger conditions. The dashed outline shows the significant clusters with higher activity in the overlap compared to the non-overlap LS conditions. (**B**) ROI-based analysis of activity in single finger and chord conditions. Error bars denote SEM across participants. ***p<0.001, two-tailed paired-samples t-test. (**C-D**) Group maps for the log-Bayes factor for the action model (C) and cue model (D). Darker colors represent stronger evidence for encoding. Each map was thresholded with PXP > 0.75 and logBF >1.

Interestingly, the areas along the intra-parietal sulcus also showed evidence for action-related encoding. Consistent with their involvement in motor planning, these areas also showed higher activation during the chords. Together with the visual areas, these areas were part of the cluster that was more highly activated during online preparation (dashed outline). Within this cluster, we found 14 cortical patches with significant action encoding and 10 cortical patches with significant cue encoding.

Outside of the area involved in online preparation, we found additional areas with strong evidence for action encoding. This was most prevalent in the PMd, which is clearly involved in motor planning before (Shenoy et al., 2013) and during (Zimnik & Churchland, 2021) motor execution. The PMd also showed much higher activity during chords than single finger presses (Figure 4B). Despite clear evidence for involvement in motor planning, we did not find any extra activation in online preparation in this area (Figure 3C, D). This suggests that the process of preparation for the next movement can run in parallel with ongoing movement control without requiring any more extra brain activity.

### Online preparation extra activation is invariant to action complexity

To investigate the processes that led to the higher activity during the overlap compared to the non-overlap condition, we asked whether the amount of extra activity differed between chords (Figure 5B) and single-finger movements (Figure 5A). If these processes were related to motor planning, we predicted that overlapping preparation would be harder if motor planning takes longer, leading to a larger overlap vs. non-overlap difference for chords. As expected, the occipito-temporal areas did not show any evidence for such a difference (Figure 5C, *F_1,21_<0.018, p>0.8951*). However, we also did not find any modulation of the overlap vs. non-overlap contrast with action complexity within the superior-parietal lobule (*F_1,21_<1.004, p>0.3884*). This suggests that the extra activity in SPL relates to processes that resolve interference associated with simultaneous action selection, independent of the exact details of the motor program.

**Figure 5.**
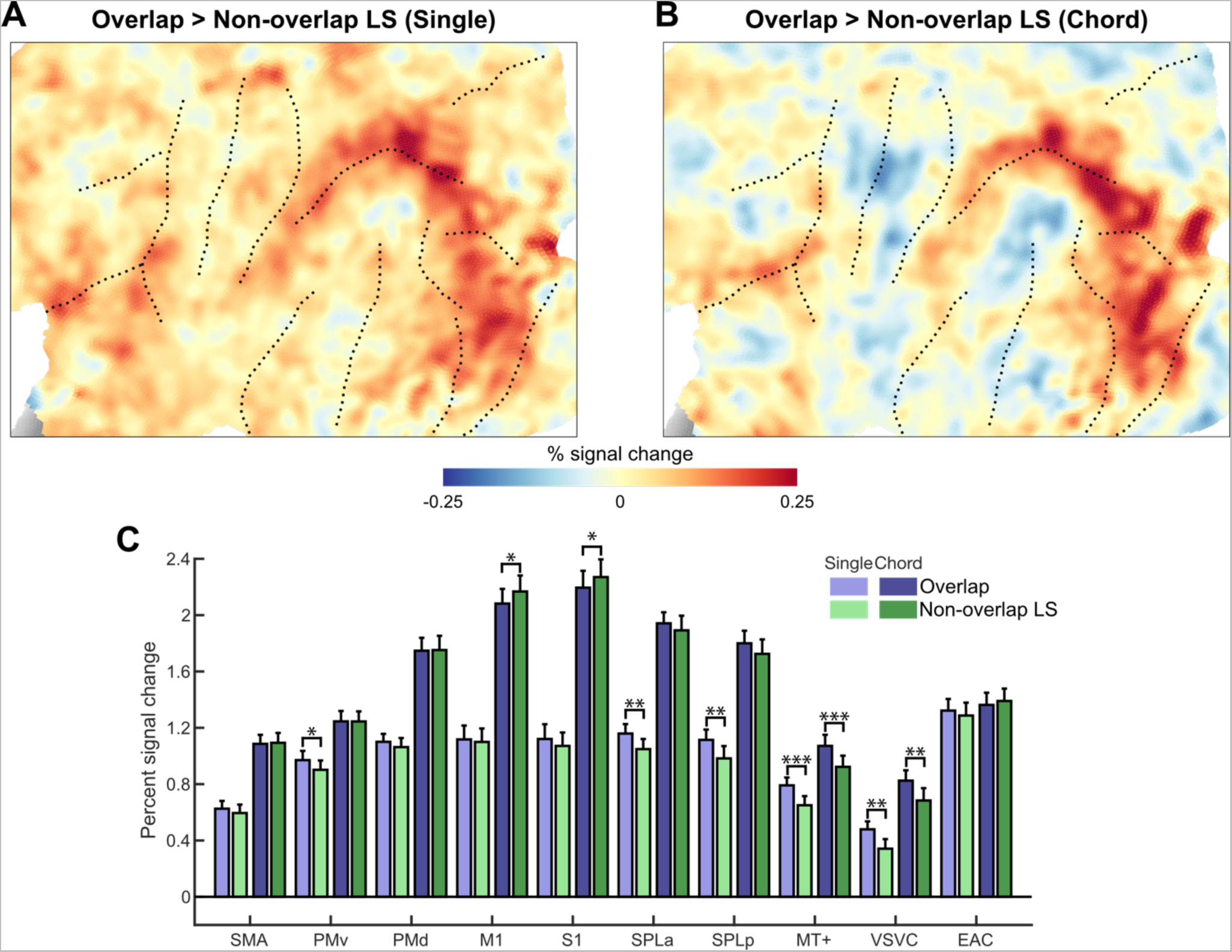
No evidence for extra online preparation activation in chord conditions. (**A**) The contrast between the overlap and non-overlap condition for single fingers and (**B**) for chords. (**C**) ROI-based analysis of percent signal change across overlap / non-overlap LS conditions for chords and single finger presses. *p<0.05, **p<0.005, ***p<0.001, two-tailed paired-samples t-test.

Taken together, our results show that only superior-parietal and occipito-temporal regions exhibit higher activation during online preparation. The extra activity was likely related to cue identification and abstract action selection but not detailed motor planning. Areas that did show clear evidence for motor planning, such as the PMd, did not show extra activity during online preparation.

## Discussion

Outside of laboratory the brain is confronted with a demanding problem: preparing the future actions while still controlling and monitoring the ongoing action. In this paper, we asked how this problem is solved at the whole-brain level. We found an unexpected dissociation: superior-parietal and occipito-temporal regions were more activated when actions overlapped, whereas premotor areas did not show any extra activity. This dissociation is all the more remarkable in that the superior-parietal lobule (SPL) and the dorsal premotor cortex (PMd) are both thought to be involved in motor planning. fMRI studies have reported that activity patterns in these two areas encode very similar information, including the intended effector (Gallivan et al., 2013; Leoné et al., 2014) and the planned sequences of future actions (Berlot et al., 2021; Gallivan et al., 2016; Yokoi & Diedrichsen, 2019). Electrophysiological recordings have also consistently shown that patterns of neural activity in both areas encode details of the upcoming movement (Cisek & Kalaska, 2005; Kalaska, 1988; Kalaska et al., 1990; Kalaska & Crammond, 1995; Kaufman et al., 2010; Nakayama et al., 2008; Scott et al., 1997; Wise et al., 1986). In our study, we found higher activity during the motorically more complex chords than during single finger presses in both areas. Furthermore, multivariate analyses showed clear evidence for action encoding, confirming the involvement of both regions in motor planning in our task. Nonetheless, we found increased activity during action overlap in SPL but not in PMd.

If online planning did occur in PMd in our task, then we need to conclude that planning can occur in parallel with an ongoing action without requiring extra activity in this area. It has been suggested that neural patterns underlying movement preparation occur in orthogonal neural dimensions to the ones supporting the motor execution (Elsayed et al., 2016; Kaufman et al., 2014). Under this arrangement, activity along the movement preparation dimensions does not interfere with the ongoing execution (Zimnik & Churchland, 2021). When the two actions overlap, the processes related to preparation and execution would also be able to superimpose in a single area without necessarily causing extra metabolic activity.

An alternative explanation for our Null finding, however, is that PMd did not perform online motor planning in our task. Although recent electrophysiological results have demonstrated that both ongoing and upcoming movements are represented at the same time in PMd (Zimnik & Churchland, 2021), these results were found in a task that involved reaches to spatial targets. In contrast, our task used an arbitrary stimulus-response mapping, which may have increased the importance of cue identification and action selection processes. It is an open question whether the direct mapping between cues and actions (Diedrichsen et al., 2001) would heighten online planning processes in PMd and may even cause increased activity during overlapping actions. However, given the clear involvement of PMd in planning actions based on arbitrary cues (di Pellegrino & Wise, 1993), we think it is more likely that online planning did indeed occur in PMd in our task, as it does during goal-directed reaching.

In contrast to PMd, we found a clear increase in activity in the overlap compared to the non-overlap condition in SPL. There is evidence that SPL is involved in motor planning (Gallivan et al., 2013; Leoné et al., 2014), but that it also represents the goal of upcoming actions independently of the exact motor requirements (Hamilton & Grafton, 2006; Henderson et al., 2022; Shushruth et al., 2022). Increased activity during the overlap condition could be attributed to interference at either stage. However, the fact that the increase was not larger in the chord than in the single finger condition seems to be at odds with interference arising at the stage of motor planning. We would have expected that when motor planning requires more time, any interference would require more activity to be resolved. Therefore, the extra activity is more likely attributed to processes preceding detailed motor planning. Neurophysiological studies have shown that SPL represents different potential actions in the form of a priority map (Bisley & Goldberg, 2010). Thus, it is possible that the selection of the two simultaneously ongoing actions causes interference, and an increase in activity is required to resolve this conflict.

Widespread increases in activity were also found in the occipito-temporal cortex. Multivariate analysis confirmed that these regions represent the visual cue but not the identity of the action. When actions overlap, the brain needs to identify the visual cue of the ongoing and next actions simultaneously. This dual task likely requires the allocation of additional attentional resources (Pashler, 1999), which then leads to increased neuronal activity. Interestingly, the activity in these visual regions was also slightly higher for the motorically more complex chords than for simple finger movements. This is surprising, as the visual and attentional requirements were tightly matched across chord and single-finger conditions. These results suggest that the identification of the visual cue and the process of motor planning do not occur in a strictly serial manner (Cisek & Kalaska, 2010): Visual areas may need to maintain the cue representation until motor planning is concluded. This would explain why complex actions that require more motor planning would be associated with higher activity in visual areas.

Taken together, our results suggest that the main bottleneck for online preparation occurs at the stage of cue identification and action selection. Cortical areas in the ventral visual stream needs to maintain a representation of the current cue while already identifying the next cue. Similarly, SPL needs to maintain the identity of the ongoing action while selecting the next action goal. These processes likely require more attentional resources when dealing with overlapping tasks (McLeod, 1977; Smith, 1968; Welford, 1952), thereby causing more neuronal activity. Conversely, the lack of extra activity in cortical premotor areas suggests that motor planning can proceed in parallel to ongoing execution, an idea that is consistent with the model of orthogonal subspaces for planning and execution (Kaufman et al., 2014; Zimnik & Churchland, 2021).

## Methods

### Participants

A total of 11 individuals (4 female, mean age = 26±4) participated in Experiment 1, and 22 individuals (12 female, mean age = 24±4) took part in Experiment 2. Four individuals participated in both experiments. Inclusion criteria required right-handedness and no prior history of psychiatric or neurological disorders. Participants provided written informed consent to all procedures and data usage before the study started, and all the experimental procedures were approved by the Human Research Ethics Board at Western University.

### Apparatus

Finger presses were produced on a right-hand MRI-compatible keyboard with five 10.5 x 2 cm keys. Each key had an indentation to guide fingertip placement. Finger presses were isometric. Forces were measured by transducers (FSG-15N1A; Sensing and Control Honeywell; the dynamic range of 0–25 N; update rate 5 ms) located beneath the fingertip indentation of each key. To register a key press, the applied force had to exceed the 0.8 N threshold, indicated by a horizontal red line on the top of the screen (Figure 1B). Five white lines were displayed on a computer screen such that the vertical position of each line was proportional to the force exerted by each finger on the respective key (Figure 1B).

### Paced response task (general procedures)

In both experiments, the task required participants to produce responses at a fixed pace. The responses were either single-finger or simultaneous three-finger presses (chord). For each of these, we assigned 5 arbitrary symbols to the 5 responses (Figure 1A). To help participants keep a regular pace, we presented a regular sequence of high and low pitch tones. Similar to the “forced-response” paradigm (Haith et al., 2016), participants had to respond synchronously with the high-pitch tones. The response time was defined as the moment summed force across fingers reached its peak.

To provide feedback on the temporal accuracy of the response, we displayed a bar in the lower part of the screen (Figure 1B). If the response was too early, the bar pointed to the left, and if the response was too late, it pointed to the right. The length of the bar indicated the size of the deviation. The acceptable deviation size was 200ms, specified by the box boundaries. If the executed response matched the instructed cue, the bar appeared in green; otherwise, it appeared in red, indicating a selection error.

Points were awarded for each press and time accuracy according to the following scheme: −2 points in case of timing error (deviation >200ms); 0 points for pressing any wrong key without timing error; 1 point in case of a correct response without timing error.

### Experiment 1

At the beginning of the experiment, participants were trained in symbol-response mapping. In separate blocks for single-finger and chord, we presented a symbol on the screen and asked them to produce the corresponding response. We encouraged them to delay their response until they felt confident that the response was correct. Each block consisted of 60 responses with symbols presented in random order, except for the fact that symbols were never repeated. We continued training until participants achieved an accuracy of above 95% for both single-finger and chord blocks.

After training, we interleaved test blocks for the two conditions. Participants produced a sequence of chords or single-finger presses at a continuous pace of 1 response per 2 seconds. A tone was presented each second, alternating between high-pitch and low-pitch sounds (gray and black notes in Figure 1B, respectively). Participants were instructed to respond simultaneously with high-pitch tones. At a random time before each high-pitch tone, a symbol appeared on the screen that instructed the required response. The preparation time for each response ranged randomly between 240 and 1750 ms. We used low-pitch tones to make response times more temporally predictable.

Five single-finger blocks were interleaved with five chord blocks. Each block consisted of 60 responses. Symbols could occur with equal probability, but no repetitions were allowed. Additionally, we switched symbol-response mapping across participants (Figure 1A).

### Experiment 2

Experiment 2 consisted of a behavioral training session and a scanning session. At the beginning of the behavioral training session, participants were familiarized with the symbol-response mapping, as done in Experiment 1. In the main task, participants again produced a sequence of responses synchronized to a sequence of regularly paced tones. These ones occurred in triplets, at a pace of 1 tone per 750ms. The first tone was low-pitch, and the next two were high-pitch, and participants were instructed to synchronize their responses with the high-pitch tones. On the screen, symbols were presented in two boxes (Figure 2A). The upper box informed participants about future responses, while the lower box contained the cue for immediately upcoming response.

In the behavioral session, we first trained participants on a simpler version of the task, giving them 1300 ms preparation time for the two responses. The symbol for each response always appeared first in the upper box 1300 ms before the upcoming high-pitch tone. 550 ms before the corresponding tone, the symbol shifted to the lower box. This transition allowed a new symbol to appear in the upper box. With this timing, the symbol for the second response emerged on the screen while the first response was still being prepared. Each pair of responses repeated 8 times (with random symbol pairs each time) – with the 16 responses forming one cycle. Before each cycle started, a 5-second instruction screen displayed either “Single” or “Chord” to indicate the press type to participants. Each training block comprised three cycles of single-finger presses and three cycles of chords. Participants completed four blocks for this phase of training.

In the main task, we trained participants to prepare for their second response in two different conditions. In the overlap condition, the preparation for the second response overlapped with the execution of the first response (Figure 2A), similar to the simplified training phase. The only difference between the overlap and training phase was that the first symbol appeared only 750ms instead of 1300 ms before the first high-pitch tone. In the non-overlap conditions, the preparation for the second response did not overlap with the execution of the first response – the stimulus for the second response appeared simultaneously with the tone for the first response. The non-overlap condition was further divided into two sub-conditions. In the non-overlap short-short (SS, Figure 2B) condition, both responses had a 750ms preparation time. In the non-overlap long-short (LS, Figure 2C) condition, the first response had a 1300ms preparation time. Trials for the 6 conditions (overlap, non-overlap SS, non-overlap LS) x (single-finger, chord) were presented in cycles of 16 responses. Prior to each cycle, a 5-second instruction screen was presented, indicating the upcoming press type (but not the overlap condition). Each block included two cycles of each condition (randomly interleaved), totalling 192 responses. Participants completed 6 blocks during training.

On the day after, participants underwent an fMRI session consisting of 10 functional runs and 1 anatomical scan. Similar to training, each block consisted of two cycles of each condition, with each cycle preceded by a 5-second instruction screen. The order of conditions was randomly interleaved. Additionally, two periods of 15 s rest, each preceded by a 5 s “fixate” screen, were placed randomly between conditions. Also, periods of 10 s rest were added at the beginning and the end of each functional run. The rest periods allowed better estimation of the baseline activation. Each of the 10 functional runs took about 6 min, and the entire scanning session (including setup and anatomical scan) lasted for about 100 min.

### Behavioral data analysis

We quantified the execution quality by evaluating the force trajectory in five-dimensional finger space from response initiation to peak force. To determine initiation time, we calculated the first derivative of the summed force (across all fingers), and then determined the moment that the rate of force change exceeded 5% of the maximum rate within that trial. If the active fingers (index and ring in Figure 1C) were pressed simultaneously and the passive fingers (like the middle finger in Figure 1C) remained stationary, the produced force trajectory (dashed lines) would be aligned with the ideal straight-line trajectory. Sequential presses of active fingers, involuntary coactivation of passive fingers, or pressing the wrong fingers all cause deviation from the ideal straight-line trajectory. We, therefore, quantified accuracy as the Euclidian norm between the produced force and the projection of the produced force onto the straight-line (orange line in Figure 1C). This distance was averaged over all time points from the initiation until peak time to produce the “mean deviation.”

We assessed the effect of preparation time on execution quality by calculating the mean deviation as a function of preparation time separately for each participant and for single-finger and chord. We then fitted the data to the following Gaussian kernel separately for each subject and each press type:

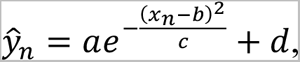

where *y_n_* is the predicted mean deviation when the preparation time is *x_n_*; a, b, c, and d are free parameters determining, respectively, the scale, the shift, steepness, and offset of the function. Parameters were then fitted to the data of press type separately using MATLAB’s fminsearch routine to minimize the mean squared error loss function. The required preparation time was estimated by equating *y_n_* to 1.05*d*.

Statistical analyses on the required preparation time and mean deviation were performed using two-tailed paired-sample t-tests and a within-subject repeated-measures ANOVA with factors conditions and press types

### Imaging data acquisition

High-field fMRI data were acquired on a 7T Siemens Magnetom MRI scanner with a 32-channel head coil at Western University (London, Ontario, Canada). The anatomical T1-weighted scan of each participant was acquired halfway through the scanning session (after the first five functional runs) using a Magnetization-Prepared Rapid Gradient Echo sequence (MPRAGE) with an isotropic voxel size of 0.75 mm (field of view = 208 × 157 × 110 mm [A–P, R–L, F–H], encoding direction coronal). To measure the blood-oxygen-level-dependent (BOLD) responses in human participants, each functional scan (352 volumes) used the following sequence parameters: GRAPPA 3, multiband acceleration factor 2, repetition time (TR) = 1.0 s, echo time (TE) = 20 ms, flip angle (FA) = 30°, 46 slices, 2.3 mm isotropic voxel size. To estimate and correct for magnetic field inhomogeneities, we also acquired a gradient echo field map (transversal orientation, the field of view: 210 × 210 × 160 mm, 64 slices, 2.5 mm thickness, TR = 475 ms, TE = 4.08 ms, FA = 35°.)

### Preprocessing and first-level analysis

Data analysis was performed using SPM12 (http://www.fil.ion.ucl.ac.uk/spm/) and custom-written MATLAB (MathWorks) routines. Images were corrected for field inhomogeneities and head motion (Hutton et al., 2002). Due to the short TR (1s), we did not adjust images for the sequence of slice acquisition. The data were high-pass filtered to remove slowly varying trends with a cutoff frequency of 1/128 Hz and coregistered to the individual anatomical scan. No smoothing or normalization to a group template was implemented during preprocessing.

The preprocessed images were analyzed with two different general linear models (GLM): The first was designed to estimate how much each of the 6 task conditions (overlap, non-overlap SS, non-overlap LS) x (single-finger, chord) activated each voxel in each of the 10 functional runs. In the design matrix, the regressor for each condition consisted of two boxcar functions (1 for each cycle of 16 responses; length 18.75 s). We also added a single regressor for instruction periods that happened before each cycle or fixation period (2 s length boxcar functions). The estimate of this regressor was not used in further analysis.

We used a second GLM to estimate the pattern corresponding to each of the 10 actions (5 single-finger presses and 5 chords, Figure 1A) within each run by modelling the first and the second response separately for each action but averaged across all responses and conditions. The regressor for each action consisted of ∼20 boxcar functions (16 responses per sequence x 2 repetitions of each sequence x 3 conditions per press type x 0.2 action occurrence probability). The length of boxcar functions varied depending on the available preparation time. We used 1.8 s for the second response in the overlap condition and the first response in the non-overlap LS condition (Figure 2A and 2C, respectively) and 1.25 s for the rest. The instruction regressor was treated the same as in the first GLM.

The boxcar functions were convolved with an individual-specific hemodynamic response function. For each participant, we tested which of 20 HRF functions drawn from the GLMsingle library (https://github.com/cvnlab/GLMsingle/) maximized the proportion of the variance that the model could explain of the time series of voxels in the left primary motor and dorsal premotor cortex. The selected HRF was then applied to the whole brain. For HRF selection, we treated all conditions as one condition; therefore, this procedure did not bias any subsequent analysis that concerned differences between conditions. Ultimately, the first-level analysis resulted in one activation image (*β*-values) per condition per run. We then calculated the percent signal change for each condition relative to the baseline activation for each voxel for each functional run and averaged it across runs.

### Surface-based analysis

Individual subject’s cortical surfaces were reconstructed using FreeSurfer (Dale et al., 1999). Individual white-gray matter and pial surfaces were extracted and spherically morphed to match a group template atlas based on the sulcal depth and local surface curvature information (Fischl et al., 1999). Subsequently, surfaces were resampled to a left-right symmetric template (fs_LR.32k.spec) included in the connectome workbench distribution. Individual data were then projected onto the group map via the individual surface.

### Regions of interest (ROIs)

We identified ten ROIs (Figure 3A) based on the cortical areas defined in Glasser et al., 2016 to cover the main anatomical areas that exhibited task-related activations in general (Figure 3B). These ROIs included the supplementary motor area (SMA), the dorsal premotor cortex (PMd), the ventral premotor cortex (PMv), the primary motor cortex (M1), the primary somatosensory cortex (S1), the anterior and posterior superior-parietal lobules (SPLa/SPLp), the MT+ complex and neighboring visual areas (MT+), the ventral stream visual cortex (VSVC), and the early auditory cortex (EAC).

The SMA was defined as the medial aspect of Brodmann area (BA) 6, covering SCEF, 6ma, and 6mp. The PMd was located at the junction between the superior frontal and precentral sulci in the lateral aspect of BA 6, covering 6a, 6d, and FEF. The PMv covered 6v, PEF, and 55b. The M1 ROI covered BA 4, cut 2 cm above and below the hand knob area (Yousry et al., 1997) to restrict it to the cortical hand area. The S1 was defined similarly as the hand-related aspect of BA 1, 2, and 3. The superior-parietal cortex was divided into an anterior region (SPLa) covering AIP, 7PC, LIPv, and LIPd, and a posterior region (SPLp) covering MIP, VIP, and 7PL. The MT+ covered areas in the lateral occipital and posterior temporal cortex, including LO1, LO2, LO3, V3CD, V4t, FST, MT, MST, and PH. The VSVC lies around the ventral aspect of the left hemisphere, covering areas anterior to early visual areas such as FFC, VVC, V8, VMV1, VMV2, VMV3, and PIT. The EAC includes A1, LBelt, MBelt, PBelt, and RI. The ROIs were defined on the group surface and then projected into the individual space via the cortical surface reconstruction of that individual. We selected all voxels that lay between the individual pial and white matter surfaces as part of the ROI.

Additionally to the ROI analysis, we also performed a continuous searchlight analysis (Oosterhof et al., 2011). A searchlight was defined for each surface node, encompassing a circular neighborhood region containing 100 voxels. The voxels for each searchlight were found in exactly the same way as for the ROI definition. As a slightly coarser alternative to searchlights, we also defined a regular tessellation of the cortical surface separated into small hexagons and extracted the functional data in the same way.

### Analysis of activation

We calculated the percent signal change for each condition relative to the baseline value for each voxel for each functional run and averaged it across runs. For the ROI analysis, the percent signal change was averaged across all voxels in the predefined regions in the native volume space of each subject. Additionally, for visualization, the volume maps were projected to the surface for each subject and averaged across the group in Workbench space.

Statistical analyses to assess differences in percent signal change were conducted using two-tailed paired-sample t-tests and within-subject repeated-measures ANOVA with factors conditions (overlap, non-overlap LS) and press types.

Statistical tests on the surface were conducted using an uncorrected threshold of *p=0.001*, and family-wise error was controlled by calculating the size of the largest superthreshold cluster across the entire cortical surface with estimated smoothness of FWHM 11.4 mm that would be expected by chance (p=0.05) using Gaussian field theory as implemented in the fmristat package (Worsley et al., 1996).

### Dissimilarities between activity patterns for responses

To evaluate which regions displayed cue- or action-specific representation, we calculated cross-validated Mahalanobis dissimilarities between the patterns of estimated activities (*β*-valus). We first pre-whitening the *β* values: We estimated the voxel noise covariance matrix from the residuals of the GLM and used optimal shrinkage toward a diagonal noise matrix following the (Ledoit & Wolf, 2003) procedure, and then divided the patterns by the matrix square-root of this estimate. Multivariate pre-whitening has been found to increase the reliability of dissimilarity estimates (Walther et al., 2016). Next, we calculated the cross-validated Mahalanobis dissimilarities (i.e., the crossnobis dissimilarities; Diedrichsen et al., 2020) between evoked regional patterns of different pairs of actions, separately for 5 single finger presses and 5 chords, resulting in a total of 2x10 dissimilarities. To obtain a measure of overall encoding, we averaged these 20 dissimilarities within each cortical surface searchlight area (Oosterhof et al., 2011).

### Model-based representational fMRI analysis

While the searchlight analysis tells us from which brain areas we can decode response identity, it does not reveal which specific aspects of the action are represented. We therefore tessellated the area with significantly positive dissimilarities using a discrete set of surface patches and then estimated the contribution of two different representational models within each patch using pattern component modeling (PCM; Diedrichsen et al., 2011). The two representational components corresponded to a cue- and motor representation, respectively. Because we do not apriori know what similarities to predict for the two components, we used the data from a group of subjects with the opposite cue-to-action assignment. We predicted that if a region represented the visual cue, the similarity between patterns should be the same as observed in the other group for the same cues (but different actions). Conversely, if a region represented the action, the pattern similarity should be the same as observed in the other group for the same action (but a different cue). Thus, we specified the two model components for group 1 from the average data from group 2 in the same area (and vice versa).

Based on these two components, we then formulated a model family containing all possible combinations of the two representational components (Yokoi & Diedrichsen, 2019). This resulted in 4 combinations, also containing the “null” model that predicated no differences among any of the activity patterns. We evaluated all 4 models using a cross-validated leave-one-subject-out scheme because different combination models had different numbers of free parameters. The component weights were fitted to maximize the likelihood of the data of N-1 subjects. We then evaluated the likelihood of the observed activity patterns for each cortical patch of the left-out subject N under that model. The resultant cross-validated likelihoods were used as an estimate of model evidence for each of the 4 models (Diedrichsen et al., 2018). The log Bayes factor BF_m_, the difference between the cross-validated log-likelihood of each model and the null model, characterizes the relative evidence for that model.

The log-Bayes factor for each model component (logBF_c_) was calculated as the log of the ratio between averaged likelihood for the models that contained the component (c = 1) versus the averaged likelihood for the models that did not (c =0) (Shen & Ma, 2019):

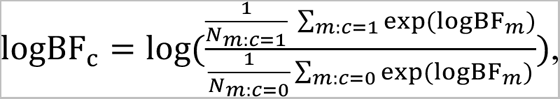

where N_m:c=1_ (N_m:c=0_) denotes the number of models (not) containing the component. Thus, a positive log-Bayes factor indicated that there was evidence for the presence of the component. The analysis was performed separately for the two groups of subjects and for single and chord data. As chord and single datasets were independent, we finally summed the log-Bayes factors for the corresponding components within each individual.

### Statistical analysis of PCM

The final log-Bayes factors for cue and action components for each participants were then submitted to a Bayesian group analysis, which estimates the probability that the component is present in a given subject (Rosa et al., 2010; Stephan et al., 2009) (spm_BMS() function implemented in the SPM 12). The significance was assessed using the protected exceedance probability (PXP) - the posterior probability that a component is present in more than half of the participants. We deemed a model contribution significant when PXP is larger than 0.75.

## Acknowledgments

This work was supported by a CIHR Project Grant to J.D. and J.A.P (PJT-175010) and the Canada First Research Excellence Fund (BrainsCAN).

## Notes

### Competing Interest Statement

The authors have declared no competing interest.

